# Identification of anthelmintic activity in a small chemical library through random screening using the worm *Caenorhabditis elegans* as a model helminth

**DOI:** 10.1101/2024.08.02.606330

**Authors:** Gemini Gajera, Vijay Kothari

## Abstract

Parasitic helminths contribute notably to the global infection burden, particularly in developing and underdeveloped nations. Anthelmintic agents currently available for therapeutic purpose are not many in number, and owing to development of resistance against them among parasitic worms, their utility spectrum is also shrinking. Hence, there is an urgent need for discovery of new anthelmintic compounds. This study subjected a small library of 181 compounds to a random screening for identification of compounds with anthelmintic activity against the model worm *Caenorhabditis elegans*. This screening effort identified 50 compounds capable of killing 100% of the worm population in less than 24 hours, at 25-50 ppm. Further investigation on these ‘hits’ is warranted to identify potential anthelmintic ‘leads’.

## Background

Helminths are parasitic worms belonging to the phyla Nematoda (roundworms) and Platyhelminthes (flatworms). They are the most common agents of infection among human populations in developing countries. Parasitic worm infections or helminthiasis affect millions of people worldwide, particularly in low-income countries with limited access to healthcare resources (Lustigman et al., 2012). The World Health Organization (WHO) recognizes helminthiasis among the neglected tropical diseases (NTD), given the limited attention and research dedicated to these infections. Moreover, many of these infections primarily affect populations in underdeveloped countries where neglected tropical diseases are prevalent. Most affected people residing in poor or middle-income countries means there is insufficient financial incentives for pharmaceutical companies to get interested in development of anthelmintic therapeutics. Effective tackling of NTD including helminth infections is considered necessary to attain the Sustainable Development Goals (WHO, 2022). In countries endemic to soil transmitted helminthiases, a third of pregnant women are infected with hookworm, which causes anaemia and blood-loss during childbirth (Aderoba et al., 2015). One example of a parasitic nematode, *Strongyloides stercoralis* infects ∼30–100 million people globally, largely in socioeconomically disadvantaged communities in the global south. Globally over 600 million people are estimated to be infected by *S. stercoralis*. The collective burden of the common helminth diseases rivals that of the high-mortality infections like HIV or malaria. According to the Technical Report of the TDR Disease Reference Group on Helminth Infections (2012), helminth infections were responsible for 85% of the NTD burden among the poorest 500 million people residing in sub-Saharan Africa. Of the 580 million people in Latin America and the Caribbean, 241 million reside in areas where at least one of the NTD is endemic. Soil-transmitted helminth infections are estimated to be borne by approximately 1.5 billion people i.e., nearly one-fifth of the world’s population. Over 260 million preschool-age children, 654 million school-age children,108 million adolescent girls and 138.8 million pregnant and lactating women are estimated to live in areas where these parasites are actively transmitted, and the need for effective treatment and preventive interventions is clearly felt (Soil-transmitted helminth infections, WHO 2023; https://www.who.int/news-room/fact-sheets/detail/soil-transmitted-helminth-infections).

Anthelmintic drugs have been instrumental in mitigating the burden of debilitating helminth infections, providing relief, and preventing severe health complications. However, the efficacy of existing anthelmintics is increasingly compromised due to the emergence of drug-resistant parasitic strains (Giunti et al., 2021). The convergence of this resistance issue with limited financial incentives for pharmaceutical companies has contributed to a disconcerting dearth in the discovery of novel anthelmintics. Parasitic worms have exhibited the capacity to develop resistance to existing anthelmintics, further limiting treatment options. Resistance of these parasites to currently used chemotherapeutic agents like albendazole and ivermectin is a matter of great concern in light of the lack of novel anthelmintics (Partridge et al., 2020; Sepúlveda-Crespo et al., 2020) in drug discovery pipeline. Repeated use of the same few anthelmintics over last 3 decades has offered enough opportunity to the parasitic populations to develop resistance against the commonly used anthelmintics.

In context of a pressing need for the discovery of new anthelmintics, preferably exerting their activity through a novel hitherto unexploited mechanism, screening libraries of natural or synthetic compounds for possible anthelmintic action can be useful in identifying potential hits. Such screening exercises need suitable model nematodes. The nematode worm *Caenorhabditis elegans* is considered to be an effective and cost-efficient model system for anthelmintic discovery (Burns et al., 2015). *C. elegans* has remained the mainstay of fundamental nematode biology research, yielding valuable information that has contributed to anthelmintic discovery and development notably (Cadd et al., 2022). Though this free-living nematode is not a parasite, it is widely believed to be a valuable model organism for anthelmintic activity assessment owing to its remarkable similarities in genetic and physiological characteristics with parasitic worms.

From literature survey (Hahnel et al., 2020; Mathew et al., 2016), it appears that the burden of nematode infections in humans, animals and plants is not negligible, and the number of anthelmintic drugs available is limited. Though *C. elegans* is not a parasitic worm, it does hold limited similarity with other parasitic worms to the extent that it can be considered a useful model for preliminary screening of anti-nematode activity (Burns et al., 2015). The present study attempted a random screening of a small library of 181 compounds present in our lab for their possible anthelmintic activity against the model worm *C. elegans*.

## Materials and Methods

### Test Compounds

We had a library of 181 compounds in our lab, which were originally procured as part of an antibacterial research project under the AIMS award programme of Atomwise Inc., USA. We attempted a blind screening approach with these compounds by randomly screening them for possible anthelmintic activity. List of all the test compounds is provided in Table S1. After procurement (from Mcule, Ukraine or Enamine, Hungary), the test compounds were immediately placed in refrigerator for storage, and reconstituted in DMSO (5 mg in 500-1000 µL; Merck) on the day of assay.

### Test Organism

Wild type N2 Bristol strain of *C. elegans* was procured from the Caenorhabditis Genetics Center (CGC, University of Minnesota). This worm was maintained on NGM agar plates (Nematode Growing Medium: 3 g/L NaCl, 2.5 g/L peptone, 1 M CaCl_2_, 1 M MgSO_4_, 5 mg/mL cholesterol, 1 M phosphate buffer of pH 6, 17 g/L agar-agar) seeded with *E. coli* OP50 (LabTIE International, Netherlands) at 22°C. For synchronization of the worm population adult worms from a 4–5 days old NGM plate were first washed with sterile distilled water, and then treated with 1 mL of bleaching solution [4% Sodium hypochlorite (Merck 61842010001730) + 1N Sodium hydroxide (HiMedia MB095-100G) + water in 1:1:3 proportion], followed by centrifugation (1500 rpm at 22°C) for 1 min. Eggs in the resultant pellet were washed multiple times with sterile M9 buffer, and then transferred onto a new NGM plate pre-seeded with *E. coli* OP50. L3-L4 stage worms appearing on this plate after 2–3 days of incubation at 22°C were used for the anti-nematode assay and were kept on NGM plates (not seeded with *E. coli* OP50) for two days, before being challenged with test compounds.

### Assay for Anti-nematode Activity

Gnotobiotic worms obtained as described above were picked from the NGM agar plate, and were distributed into different wells of a 24-well plate (HiMedia) containing M9 buffer. Ten hermaphrodite worms were delivered per well. This was followed by addition of required volume (5 or 10 µL) of the DMSO-dissolved test compound so as to achieve final concentration of 25 or 50 ppm. Total volume in each well was kept 1 mL. Three replicates for each concentration were set. Control wells contained worms in M9 buffer (with no test compound). Benzimidazole (HiMedia) and ivermectin (SRL) were used as positive controls. Since these compounds were dissolved in DMSO (Merck), an appropriate vehicle control (i.e. worms in M9 buffer + 0.5%v/v DMSO) was also set. These 24-well plates were incubated at 22°C for 5-days, and a live-dead count was performed under microscope (4X) on daily basis. Non-moving straight worms were considered as dead. Plates were tapped to confirm lack of movement in dead-looking worms. On last day of experiment, when plates could be opened, dead-looking worms were touched gently with a straight wire to further confirm lack of response.

### Statistics

All values reported are means of three experimental replicates, and measurements are reported as mean ± standard deviation (SD). Statistical significance of the data was evaluated by applying t-test using Microsoft Excel^®^. *P*-values ≤0.05 were considered to be statistically significant. Compounds showing potent anthelmintic activity were subjected to one more independent assay (comprising three replicates) for confirming reproducibility.

## Results

Of the 181 test compounds, 18 were insoluble in DMSO. Of the remaining 163 DMSO-soluble compounds, 58 were able to kill ≥50% of the worms within 72 hours. Of these 58 compounds (Table 1), 50 were able to kill 100% worms in less than 24 hours. Since we had only limited quantity of the test quantity available, we could test each of them either at single (25 ppm) or two different concentrations (25 ppm and 50 ppm). While the active compounds identified in this study showed activity better than one of the positive controls, benzimidazole, their direct comparison with another positive control ivermectin is possible only by testing these compounds at a concentration as low as 1 ppm at which ivermectin could kill 100% worms in less than 24 hours. While all active compounds are listed in Table-1, a representative survival plot showing results for some of the test compounds and both positive controls is presented in Figure-1.

**Table 1.**
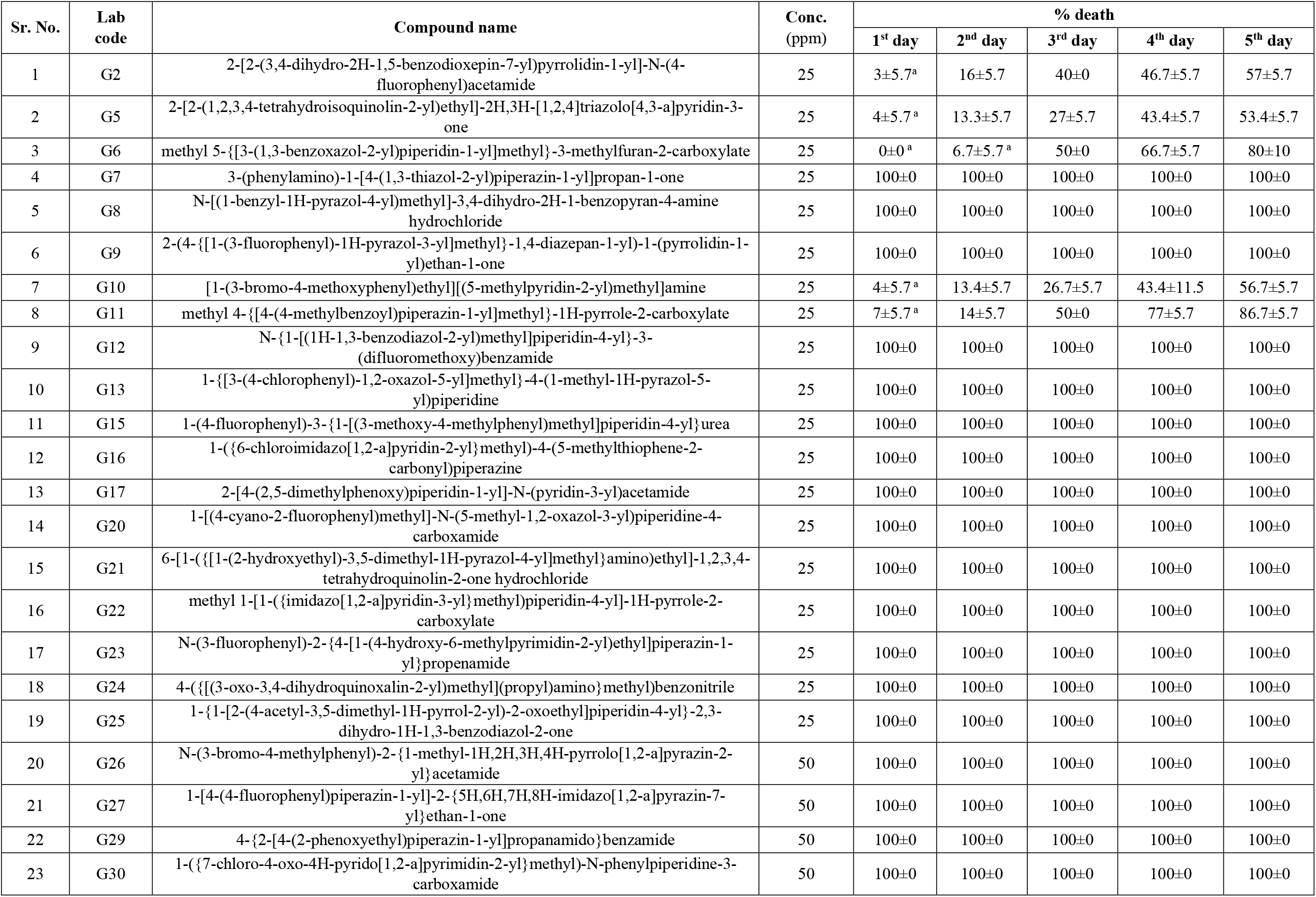
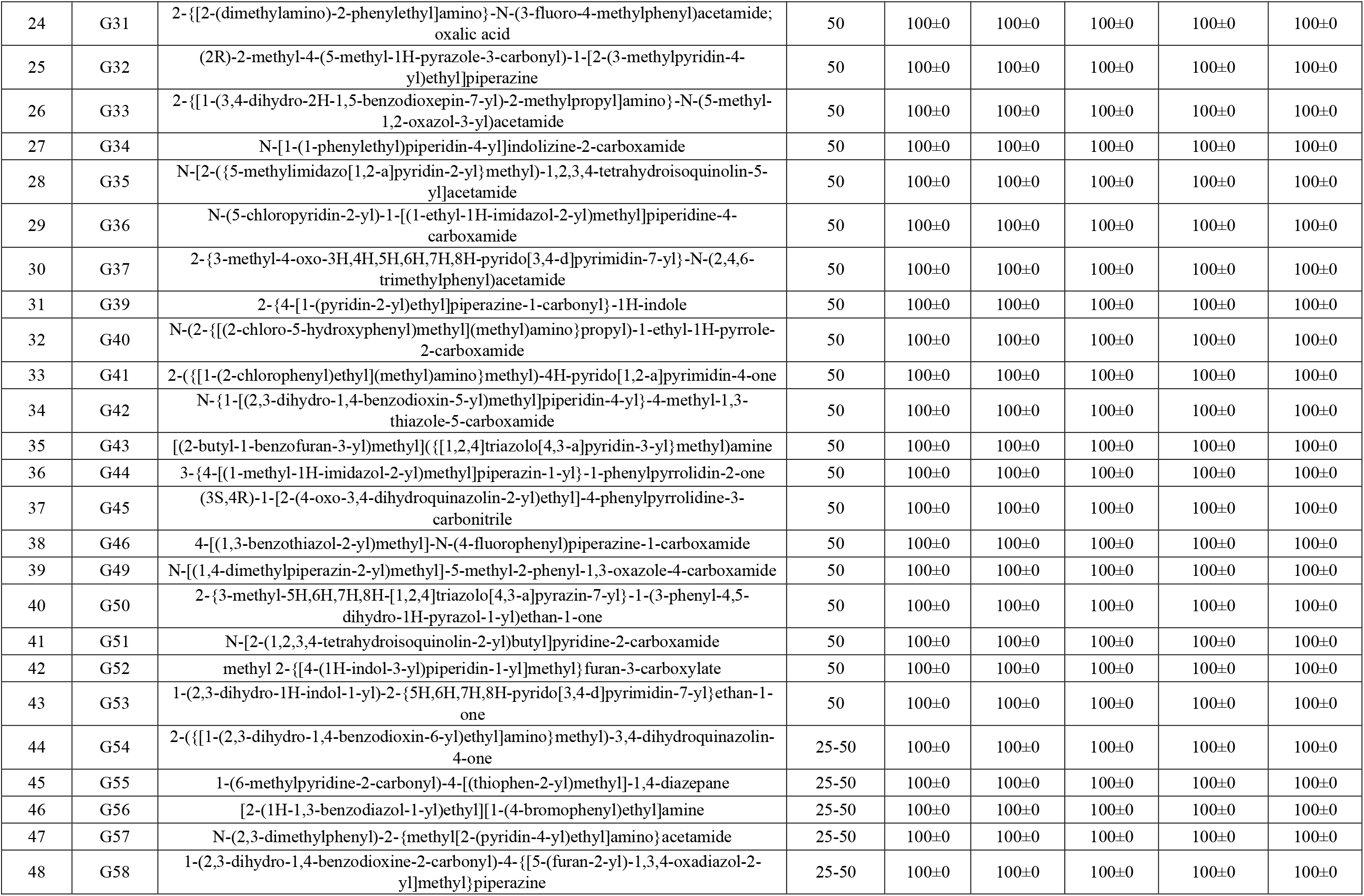
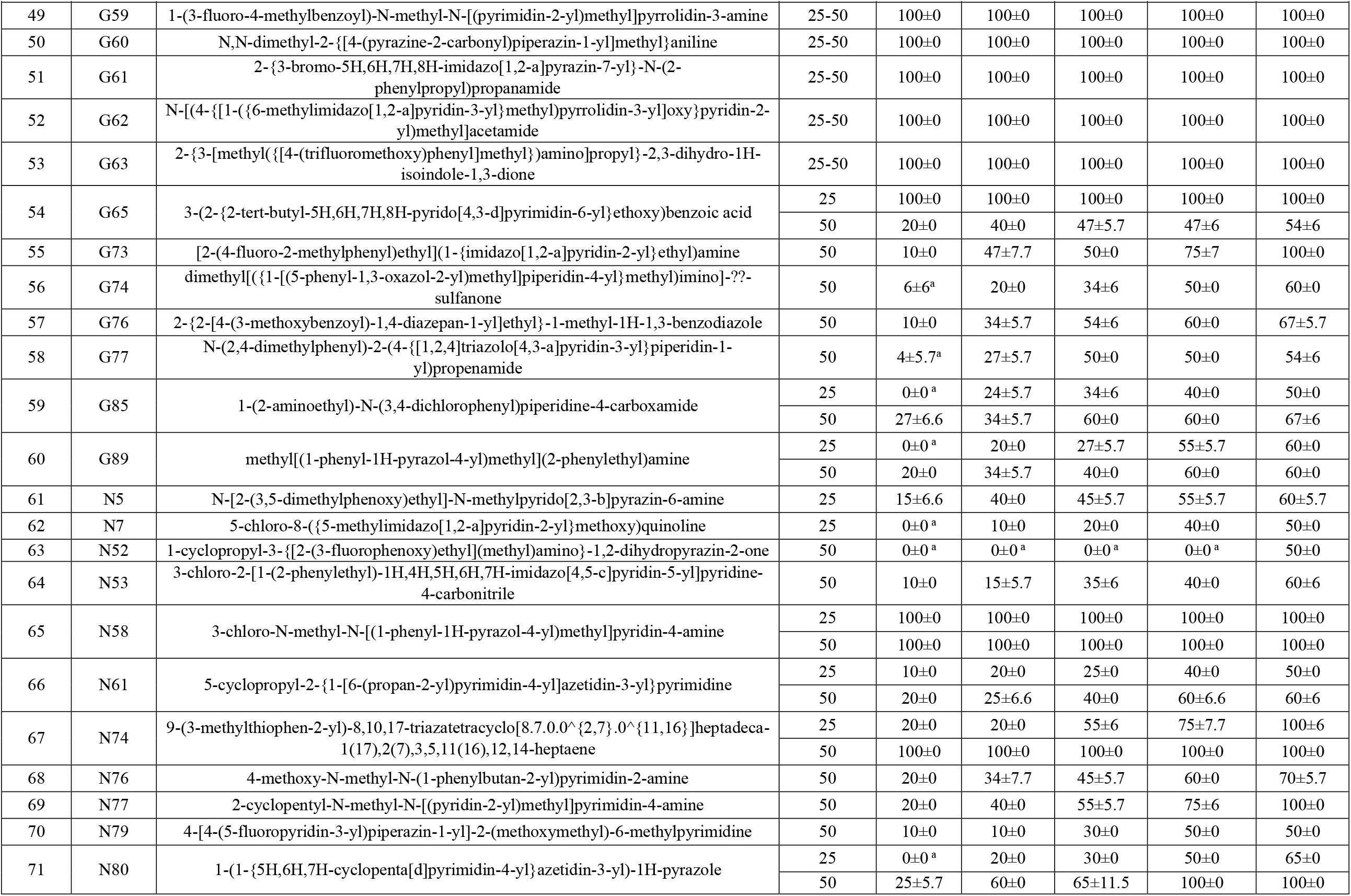
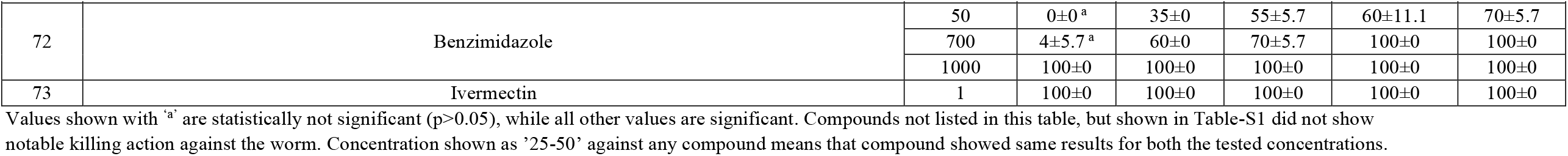
Compounds identified to possess anthelmintic activity against *C. elegans*.

**Figure 1.**
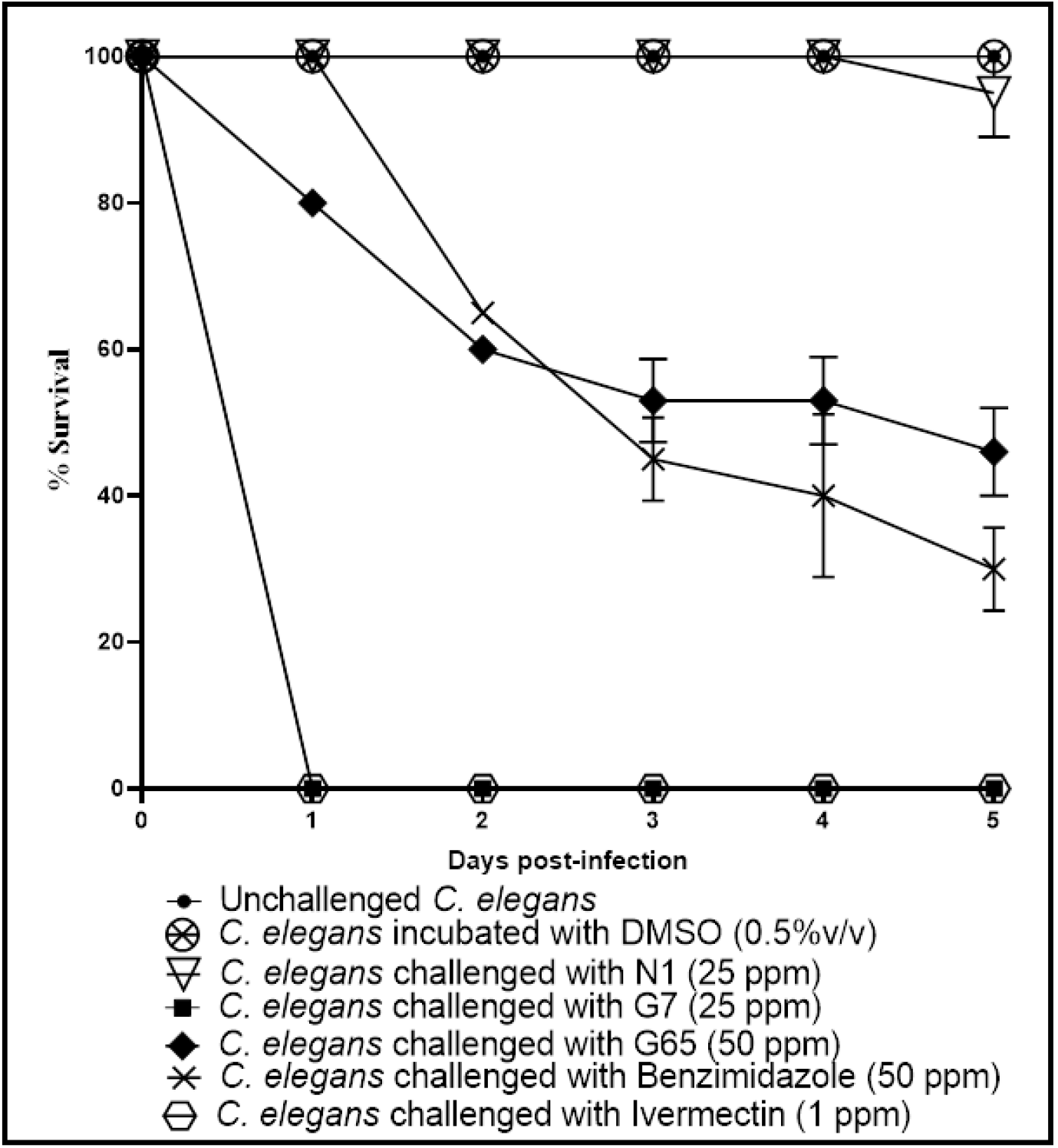
A representative survival plot depicting anthelmintic activity of some test compounds and that of positive controls. For the purpose of this plot, we have selected one compound with no anthelmintic activity (N1), one capable of killing ∼50% worms at a slow pace (G65), and one capable of killing all worms within a day (G7). Benzimidazole was tried at 700 ppm too, wherein it could kill all worms over a 4-day period, and at 1000 ppm it could kill all worms within a day.

At the time of writing this manuscript, our search on PubChem for the compounds identified as possessing anthelmintic activity in this study indicates that this is the first report showing any kind of biological activity in these compounds. The top fifty anthelmintic compounds identified in this study should be tested at still lesser concentrations (preferably against some human-parasitic nematode) to find out the most potent anti-nematode candidates from among them, and ‘time required to kill’ assay should also be done for them. Safety of the anti-nematode compounds identified in this study for higher animals/humans also needs to be explored. Molecular mechanisms associated with the anti-nematode activity of the most potent compounds should be elucidated. While it is beyond our expertise, it will be interesting and useful to see whether there are any structural similarities among some of the potent anthelmintic compounds reported in this study.

## Acknowledgement

Authors thank Nirma Education and Research Foundation (NERF), Ahmedabad, for infrastructural support; and Atomwise Inc., USA for making the test compounds available. GG acknowledges scholarship from the Government of Gujarat through their SHODH scheme.

